# WAS IT A MATch I SAW? Approximate palindromes lead to overstated false match rates in benchmarks using reversed sequences

**DOI:** 10.1101/2023.06.19.545636

**Authors:** George Glidden-Handgis, Travis J. Wheeler

## Abstract

**Background:** Software for labeling biological sequences typically produces a theory-based statistic for each match (the E-value) that indicates the likelihood of seeing that match’s score by chance. E-values accurately predict false match rate for comparisons of random (shuffled) sequences, and thus provide a reasoned mechanism for setting score thresholds that enable high sensitivity with low expected false match rate. This threshold-setting strategy is challenged by real biological sequences, which contain regions of local repetition and low sequence complexity that cause excess matches between non-homologous sequences. Knowing this, tool developers often develop benchmarks that use realistic-seeming decoy sequences to explore empirical tradeoffs between sensitivity and false match rate. A recent trend has been to employ reversed biological sequences as realistic decoys, because these preserve the distribution of letters and the existence of local repeats, while disrupting the original sequence’s functional properties. However, we and others have observed that sequences appear to produce high scoring alignments to their reversals with surprising frequency, leading to overstatement of false match risk that may negatively effect downstream analysis.

**Results:** We demonstrate that an alignment between a sequence S and its (possibly mutated) reversal tends to produce higher scores than alignment between truly unrelated sequences, even when S is a shuffled string with no notable repetitive or low-complexity regions. This phenomenon is due to the unintuitive fact that (even randomly shuffled) sequences contain palindromes that are on average longer than the longest common substrings shared between permuted variants of the same sequence. Though the expected palindrome length is only slightly larger than the expected longest common substring, the distribution of alignment scores involving reversed sequences is strongly right-shifted, leading to greatly increased frequency of high-scoring alignments to reversed sequences.

**Impact:** Overestimates of false match risk can motivate unnecessarily high score thresholds, leading to potentially reduced true match sensitivity. Also, when tool sensitivity is only reported up to the score of the first matched decoy sequence, a large decoy set consisting of reversed sequences can obscure sensitivity differences between tools. As a result of these observations, we advise that reversed biological sequences be used as decoys only when care is taken to remove positive matches in the original (un-reversed) sequences, or when overstatement of false labeling is not a concern. Though the primary focus of the analysis is on sequence annotation, we also demonstrate that the prevalence of internal palindromes may lead to an overstatement of the rate of false labels in protein identification with mass spectrometry.

## Introduction

The utility of tools that label biological sequences is dependent on the ability to reasonably predict the risk that a proposed label is due to some random or biased process, rather than true signal. An example is the case of sequence homology detection, in which an unlabeled sequence *q* is compared to a collection of known sequences, and similarity to one or more of those sequences serves as the basis of annotation of *q*.

Homology detection tools typically represent the risk of false annotation by providing the supporting alignment’s E-value along with its score *S*. For search of a particular query sequence against a given target database *T*, the E-value gives the number of alignments expected to meet or exceed the score *S* if *T* consisted exclusively of unrelated (random) sequences. Theoretically-grounded calculation of the E-value has been established for gap-free alignments (1, 2), and empirical studies demonstrate that these E-value predictions are accurate for search against randomly generated sequences of nucleotides or amino acids (3, 4).

Even so, genomics practitioners are keenly aware that these E-values are often overestimates of the confidence with which we should label one sequence as being related to another, so that it is common for analysis pipelines to require seemingly-extreme E-value cutoffs for search matches (5). This is because the models underlying the scoring schemes used in alignment (6, 7) assume that the database sequences are generated under a simple random distribution (sample with replacement), whereas true biological sequences are not always random in this way.

One notable deviation comes in the form of locally repetitive sequence. So-called tandem repeats are typically the result of some form of replication error (8), and can present with a repeat period ranging from 1 to several hundred (9). They produce substantial risk of spurious matches when both query and target sequence contain (possibly degenerate) repeats with similar repeated units. One strategy for compensating for tandem repeats is to employ a tandem repeat *masking* tool (10–12) to hide regions of repetitive sequence away from homology detection. Such strategies are necessarily limited by the selection of some score threshold, and may thus either over-mask (hiding true homologs) or under-mask (allowing decayed repetitive regions to be identified as true homologs (13)).

Continuing concerns about real-world accuracy of E-values, and of the compensatory score adjustment/masking procedures, have lead researchers to evaluate tools with benchmarks that *empirically* measure false positive annotations in addition to true positive recall, rather than rely solely on theoretical E-values. A benchmark will generally contain a set of *query* sequences and a distinct set of *target* sequences. In the target set, some sequences are related to a query (these are the *positives* that the tool will ideally annotate as such), while other sequences are not related to a query (these are *negatives* or *decoys* that the tool should not annotate). The benchmarked algorithm is used to compute alignments without knowledge of true labels, and a predicted classification is produced. Wherever the computed classification agrees with ground truth, the algorithm is said to have produced a *true positive* or *true negative*, and elsewhere it produces *false negatives* and *false positives*, respectively.

A variety of techniques have been developed to automate the production of positive and negative sequence pairs. In this paper, we are particularly interested in methods of creating negative sequence pairs, because these decoys inform efforts to establish score cutoffs that limit false discovery rate. One option for generating decoy sequences is to randomly sample real biological sequences from a large dataset. This strategy is likely to produce an overestimate of the false positive rate because each randomly chosen ‘decoy’ has some possibility of being homologous to the query. Alternative methods for producing decoys include (i) shuffling the letters of a real biological sequence, or (ii) generating a sequence by sampling letters under a random model. In both of these cases, failure to include repetition or other realistic features of biological sequences may lead to an underestimate of the rate of false positives that will be observed when the tool is applied to real biological sequence.

Another mechanism for producing non-homologous bio-realistic sequence is to use reversed biological sequences, so that decoys are generated by selecting true biological se-quences and reversing the order of their characters. Reversal has several immediate benefits over stochastic methods. Reversal preserves the distribution of letters in a sequence, and also preserves the existence of short (possibly degenerate) tandemly repetitive regions and regions of low compositional complexity that are often found in biological sequences. Meanwhile, the reversed sequence almost certainly does not fold or behave the same as the original.

Thus, a decoy set consisting of reversed sequences is expected to possess the low complexity and repetitiveness of real biological sequence, but with no true homology to an actual sequence. Such sequences demonstrate much higher rates of apparent “false” matching than do shuffled sequences, and it is tempting to view these as a better (more realistic) benchmark for false labeling, in the context of both nucleotide (11) and protein (14) annotation. We have employed reversed sequences in this way (15) for the same reasons; however, during post-publication analysis of our benchmark results, we realized that “false” matches of transposable element queries to a reversed (not complemented) genome were surprisingly likely to be to “self” – a match of a query sequence to the reversed genome was highly likely to be to a reversal of an instance its own family. In an earlier paper evaluating software for protein search, Eddy (16) suggested the source of this observation, while explaining why that manuscript’s analysis did not include reversed decoy: (“reversed sequences are surprisingly significantly more likely to show a significant match to the original sequence […] because of a counterintuitive statistical effect of the frequency of approximate palindromes”). Here, and throughout this manuscript, the term “palindrome” follows the canonical definition and refers to a sequence that reads the same backwards and forwards (e.g. 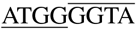 or 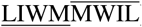). This is in contrast to an alternate usage of the term “palindrome” that sometimes appears in genomics literature: a sequence that consists of two spaced or adjacent inverted repeats, in which matching sequences occur as reverse complement (e.g. 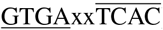).

Here, we explore the phenomenon of surprisingly high palindrome lengths, and the extent to which they may cause the use of reversed decoys to overstate the risk of false discovery in sequence database search tools. Through empirical analysis, we demonstrate that this palindromic phenomenon explains much of the increase in false discovery relative to shuffled sequence.

### Why are unexpectedly high scores for reversed sequences a problem?

It has recently become common to present the accuracy of homology search tools by reporting the percent of all true instances that are assigned an E-value better than the highest-scoring decoy (a measure that we call “recall before first false positive”, RBFFP, as used in (14) and (17)). When the top-scoring decoy produces E-value *D*, RBFFP will ignore differences in sensitivity that occur for matches with E-value *> D*. If the method of generating decoys is prone to producing false matches with inappropriately high score (low E-value), then the RBFFP-imposed cutoff for differentiating between tools will be extreme, such that sensitivity differences in interesting score ranges are ignored. For example, a decoy with *D*=1e-10 will ensure that RBFFP does not distinguish between one tool that reports 20 matches with E-value between 1e-5 and 1e-10, and a separate tool that only reports a single match in that range. Failure to distinguish tools producing different rates of recall in this ‘marginal’ score range may be concerning, particularly when there is a risk that the ‘marginal’ cutoff might be unintentionally set higher than is appropriate.

As we demonstrate in the Results section, reversed decoy sequences are prone to producing surprisingly high scores (low E-values). As a result, we suggest that care be taken in using reversed sequences as decoy sequences when evaluating the occurrence of false discovery (14) or selecting score thresholds for reporting (13). If reversed sequences are to be used as decoys, the input sequences should first be filtered to remove matches to the query set. Alternatively it may be preferable to use simulated sequences in which care is taken to produce realistic sequence (for DNA, see (18)). We hope that this manuscript will induce exploration into alternative mechanisms for improving the generation of bio-realistic decoy sequences (particularly proteins) for approximating false similarity rates.

### Intuition and Background for Palindrome Frequencies

We find it instructive to consider a simple calculation related to the expected frequency of a length-*n* palindrome in a string *A*, and of a length-*n* common substring in two strings *A* and *B*, both over a uniformly distributed alphabet of size *k*. The next two paragraphs are a sketch, intended to provide intuition about expected palindrome length; they do not represent a complete analysis.

Consider some position *i* in *A*. Ignoring edge cases, the chance that this is the center position of a length-*n* palindrome (where *n* is odd) is (1*/k*)^(*n*−1)*/*2^ – a length-*n* palindrome is formed if the (*n*−1)*/*2 letters to the right if *i* are matched (in reverse) to the (*n*−1)*/*2 letters to the left of *i*, each with 1*/k* probability. This corresponds directly to the probability that position *i* in A and position (*m*−*i* + 1) in the reversal of A (which are the same letter) will be the center of an identical common substring of length *n*.

Contrast this with the chance of observing a length-*n* identical common substring in two length-*m* strings, *A* and *B*. For some position *i* in *A* with value *a*_*i*_, with reasonably large string length *m* and modest value *k*, there will almost certainly be some position *j* in *B* with value *b*_*j*_=*a*_*i*_; this is a length-1 common substring between *A* and *B*. Ignoring edge effects, the chance that this is the center position of a length-*n* common substring is (1*/k*)^(*n*−1)^, because each of the neighboring *n*−1 characters in *A* has a 1*/k* chance of being matched by the corresponding position in *B*. For each position *i*, there are expected to be *m/k* candidate centers for common substrings, but this number is dwarfed by the much lower probability of producing a common substring of even modest length.

The above sketch ignores edge effects and cases of even-length or approximate palindromes, yet suggests that with a randomly sampled sequence *A*, a length-*n* palindrome is less surprising than is a length-*n* substring shared by *A* and some other randomly sampled sequence *B* (or a shuffled variant of *A*). Generally, the expected length of the longest common sequence (LCS) of two strings of length-*k* is 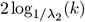(19), and the expected longest palindrome length in a length-*k* string is 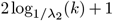. (The probability amplitude *λ*_2_ is determined from frequencies of each letter; for an alphabet of size *s*, on uniformly-distributed sequences, *λ*_2_ = 1*/s*).

Finally, we consider approximate palindromes. An *α*-gapped palindrome is a kind of approximate palindrome defined as *w* = *uvū* where *u* and *v* can be any strings and *ū* is the reversal of *u*. We say *w* is *α*-gapped whenever 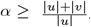. Duchon et al. (20) give an expectation for the length of *α*-gapped palindromes in a length-*k* string (*L*_*k*_) as 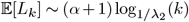 or equivalently 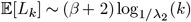 where 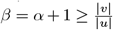. By considering odd-length palindromes as gapped palindromes with |*v*| = 1, it follows that 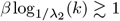, and it can be demonstrated that the expected length of odd maximal palindromes becomes 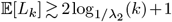 [manuscript in prep]. To our knowledge, there is no further published analysis of approximate palindromes involving substitutions or a more general pattern of gaps distributed throughout the palindrome.

Though the expected (mean) palindrome length of a sequence is only slightly higher than the expected length of the longest common substring of that sequence and its shuffled partner, we hasten to note that exact palindromes are not exclusively responsible for high scoring alignments; they simply provide a seed from which high-scoring alignments can expand, and that expansion follows mirrored match scoring dynamics that are similar to the creation of palindromes but sufficiently complicated to exceed the scope of this manuscript. We also highlight that the mean match length does not reflect the expected frequency of outliers, which will have almost no effect on the mean due to their rarity; it is these rare high-scoring matches that drive most of the alignment score results seen below.

### Paper Outline

In the following sections, we explore the connection between approximate palindrome length and the increased scores produced when aligning sequences against their reversal, relative to aligning against shuffled sequences. We begin by describing the sequence data used in our analyses, then examine the extent to which sequences achieve higher than random score when aligned to their reversal. We follow this with a study of the frequency of long palindromes and common substrings, and an investigation of the the roles of tandem repeats and mutations in the context of using reversed sequences as decoy sequences. Finally, we explore the impact of palindromes on assessing the false discovery rate in sequence annotation and in mass spectrometry analysis.

## Methods

### Preparation of Datasets

#### Real protein sequences and their paired relatives

To gain insight into the influence of decoy type on the distribution of high-scoring matches between sequences, we developed related sets of paired protein sequences. For each pair, we first identified one sequence *q* from Swiss-Prot (21), then either (a) selected another Swiss-Prot sequence *t*, (b) shuffled *q*, or (c) reversed *q*.

It is possible that two sequences that are randomly selected from Swiss-Prot will in fact be homologous.To reduce this risk, we ensured that paired sequences did not share folds as defined by the SCOP database (22, 23). Specifically, sequences from release 2022_04 of Swiss-Prot were length-filtered to identify 565,854 sequences with length between 50 and 2000; this set is called **sprot**_**all**_. These were aligned to all SCOP2 superfamily representatives using blastp from the BLAST suite (version 2.12.0+) (24). Swiss-Prot sequences were retained for this dataset only if they yielded an alignment that covered at least 95% of the length of both the Swiss-Prot and SCOP sequence – this ensured that we could reasonably annotate all folds on retained proteins. The result is a set of 106,740 sequences, called (**sprot**). Length distributions are shown in Figure 1.

**Fig. 1.**
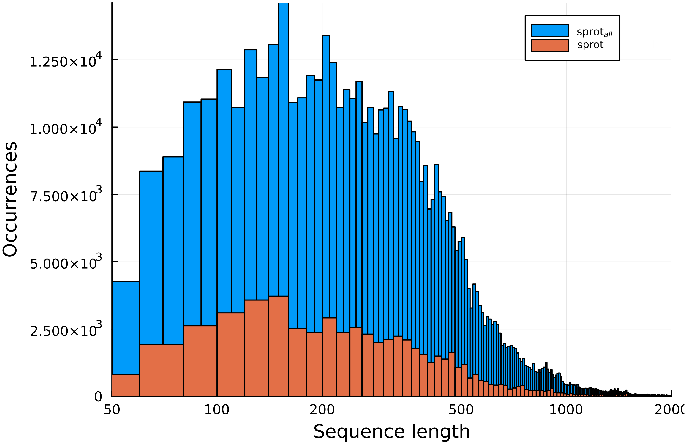
Distribution of sequence lengths for sequences sampled from Swiss-Prot (**sprot**_**all**_) and for those that match a SCOP family (**sprot**).

Sequences in (**sprot**) were assigned to clusters according to their SCOP superfamily. For each sequence *q* in **sprot**, its cluster *v* was identified, and a distinct cluster *w* ≠ *v* was chosen via sampling weighted by cluster size. A sequence *t* was then sampled from *w* and assigned to **random**. This second set (**random**) contains presumably non-homologous partners for the set **sprot**, and can be used to produce an alignment score distribution by aligning the *i*^*th*^ sequence of **sprot** to the *i*^*th*^ sequence of **random**.

To explore the distribution of scores when aligning a biological sequence *q* to a shuffled copy of *q*, each sequence in **sprot** was shuffled to produce the set **shuf**; the *i*^*th*^ sequence in **shuf** will share length and amino acid composition with the *i*^*th*^ sequence in **sprot**, but no homology. Similarly, each sequence in **sprot** was reversed to produce the set **rev** (note: this is true reversal, not reverse complementation). All shuffling and reversals were computed using the esl_shuffle tool from the Easel package as released in HMMER 3.3.2 (25).

It is plausible that the high rate of palindromes in biological sequences is due to repetition in those sequences. To demonstrate that the palindrome phenomenon is general to all sequences, and not just repetitive ones, we reversed the sequences of **shuf** to produce **shufrev**; because the shuffled sequences do not retain signal of repetition found in true proteins, an alignment of a sequence in **shuf** to its mate in **shufrev** will highlight pure features of reversal alignment. Analogous sets of unrestricted relatives of **sprot**_**all**_ are called **rev**_**all**_, **shuf**_**all**_, and **shufrev**_**all**_ respectively. Figure 2 provides a visual guide to the various alignment pairings used through-out the manuscript.

**Fig. 2.**
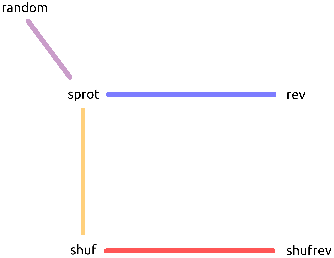
The core experiment design compares five related sequence sets, **random, sprot, rev, shuf**, and **shufrev**, represented here as nodes, by means of pairwise alignment. Each colored edge represents a comparison of two sequence sets. For each connected pairing, the edge color corresponds to the color used for that pairing throughout the manuscript in both figures and text.

#### Reversals in mutated sequences

In the reversal analyses described above, each **sprot** sequence is compared to a reversal of itself. In a carefully constructed benchmark, query sequences themselves may not be present among the sequences that are reversed to produce the decoy set, calling the applicability of this analysis into question. But the benchmark may contain their homologs, which will show some level of sequence similarity to the queries. To understand how the level of divergence impacts distribution of scores induced by approximate palindromes, we generated six mutated variants of **shuf**_**all**_, and reversed each to form a new diverged reversed benchmark. For each variant, a conditional distribution was used as the basis of mutation, so that *s* = *s*_1_…*s*_*n*_ in **shuf**_**all**_ was transformed to a related sequence *t* = *t*_1_…*t*_*n*_, in which *t*_*i*_ is an amino acid sampled from the distribution *p*(*x*|*s*_*i*_). For the first transformation, substitution probabilities were derived (26) from the BLOSUM90 substitution matrix. This matrix has a mean self-emission frequency (*p*(*a*|*a*)) of 45.6%. Additional transformations were produced based on modified distributions in which the likelihood of mutation was reduced for each amino acid *a*, adjusting the likelihoods of mutation to the other 19 amino acids *p*(*b*|*a*) proportionately. These distributions have average self-emission frequencies (*p*(*a*|*a*)) of 50%, 60%, 70%, 80% and 90% respectively. These enable consideration of the likelihood of finding matches between a query and reversals of homologs that show varying levels of percent identity to the query. These augmented distributions preserve the relative frequency of mutations derived from BLOSUM90 (*p*(*b*|*a*) for *b ≠a*) while increasing the expected percent identity between the original sequence *s* and its relative *t*. We refer to these distributions as **shufrev**_**all**_**-46pct, shufrev**_**all**_**-50pct, shufrev**_**all**_**-60pct, shufrev**_**all**_**-70pct, shufrev**_**all**_**-80pct**, and **shufrev**_**all**_**-90pct**, respectively.

#### Repeat Masking

To further investigate the role of repetition in false matching for biological sequences, we hard-masked repetitive sequences in **sprot**_**all**_ using *tantan* version 40 (11) with default parameters, producing **masked sprot**_**all**_.

#### Simulated fragmentation with trypsin

In mass spectrometry-based proteomics, proteins are broken into fragments, often employing trypsin digestion. The identity of the resulting fragments is determined based in part on their mass. Typically, the fragments evaluated by mass spectrometry have length 5 to 100 amino acids because the signal induced by short sequences (*<* 5 residues) is indistinguishable from noise, while long sequences (*>* 100 residues) may exceed the the physical limits of the machine. To simulate this process, we applied a simplified approximation of trypsin digestion as follows. Consider a protein *P* = (*p*_1_, *p*_2_, …, *p*_*n*_) and its set *S* = (*s*_1_, *s*_2_, …, *s*_*k*_) of potential trypsin digestion positions at which there is an arginine or lysine, ignoring any position that is not followed by a proline. Taking a pair of positions (*s*_*i*_, *s*_*j*_) from *S, P* may be split into three fragments, (*p*_*1*_..*p*_*si*_), (*p*_*si*_+1..*p*_*sj*_), (*p*_*sj*_ +1..*p*_*n*_). For all protein sequences in **sprot**_**all**_, we enumerated all pairs (*i, j*) of potential digestion sites and produced all resulting three-way splits, then filtered to remove fragments with length below 5 or greater than 100. The remaining fragments were accumulated into the set **sprot**_**tr**_. Each member of **sprot**_**tr**_ was shuffled to produce a shuffled control denoted **shuf**_**tr**_. This approach does not capture the true complexity of trypsin digestion patterns (27), but is sufficient for our purposes of analyzing palindromes in relatively short fragments.

#### Annotation of Chromosome 22

The preceding datasets only comprise protein and peptide sequences. To demonstrate that the elevated presence of approximate palindromes in sequence benchmarks is not merely a feature of amino acid sequences, we consider DNA sequences from the human genome.

The sequence of human chromosome 22 in hg38.p14 along with the NCBI RefSeq annotation was downloaded from UCSC’s Genome Browser (28). The command-line tool GffRead (29) was used to extract and concatenate exon regions into sequences corresponding to the mRNA transcripts of chromosome 22, denoted **mRNA**. The intron and intergenic regions of chromosome 22 were also sampled for sets of sequences with similar lengths as **mRNA**, denoted **intron** and **intergenic**.

As above, shuffling and masking were performed, and the resulting variant datasets are denoted **shuf**_**mRNA**_, **shuf**_**intron**_, **shuf**_**intergenic**_, and **masked mRNA, masked intron**, and **masked intergenic**.

### Software and algorithms

Exploration of score and count distributions was achieved using established tools and algorithms. For the large Swiss-Prot-based dataset, sequence database search scores were computed using *blastp* (24). For our purposes, pairwise alignment was performed by creating a blast database for each singular target, and this target was “searched” with its paired query sequence. The parameter for (minimum) word length was set to 4. The default substitution matrix (BLO-SUM62, half-bit) and gap penalties (gap open = −11, gap extend = −1) were left unchanged. One *blastp* alignment may contain several results, each with its own score – for our experiments, only the highest-scoring match was retained and reported. We report score, rather than the associated E-value, because E-value is dependent on the size of the query and target datasets, while score is stable for an aligned pair.

One problem with the BLAST-based analysis is that for many comparisons, the similarity between sequences is sufficiently poor that not even a low-scoring *blastp* result is produced. To gain insight to the distribution of lower-scoring results, we used the BioJulia implementation of Smith-Waterman (SW) algorithm BioAlignments.jl (30) for finding the optimal local alignment between two sequences. The algorithm is parameterized by a substitution matrix (whose values at row *i* and column *j* determine the cost of substituting letter *i* with letter *j*), as well as gap-open and gap-extend penalties, which determine the cost of skipping one or more characters in either sequence. When gap penalties and off-diagonal values in a substitution matrix are very large negative numbers (e.g., −1*e*12), Smith-Waterman will never produce gaps or mismatches and effectively performs exact alignment; we use this mode for computing longest common substrings (LCS) between sequences. In all other cases, the gap penalties and substitution matrix are set to BLAST’s defaults.

Using either BLAST or Smith-Waterman (with typical parameters) for reversed alignment readily produces approximate palindromes, but rarely identifies true palindromes. To measure the frequency of exact palindromes in Swiss-Prot, we implemented Manacher’s algorithm (31) in Julia.

Code used to produce all results and figures may be found in Supplementary Material at https://github.com/TravisWheelerLab/WASITAMATchISAW.

## Results

### Alignment scores for Swiss-Prot variations

To understand the influence of reversals on sequence alignment score distributions, we aligned sequences taken from Swiss-Prot (length between 50 and 2000) against a variety of decoy datasets. Each pairing involves either 106,740 (**sprot** and relatives) or 565,855 (**sprot**_**all**_ and relatives) pairwise alignments, with the *i*^*th*^ sequence of one set aligned to the *i*^*th*^ sequence of the other set using *blastp* (see Methods for definitions of sequence sets). The first pairing (**sprot**↔**random**) replicates the case in which a real protein sequence is aligned to some randomly chosen (presumed non-homologous) protein. The second pairing (**sprot**↔**shuf**) corresponds to a simple decoy strategy: the true protein is aligned to a sequence that is a shuffled copy of the original – it shares length and amino acid composition, but is not a true protein. As seen in Figure 3A, the distribution of scores in these two pairings are similar.

**Fig. 3.**
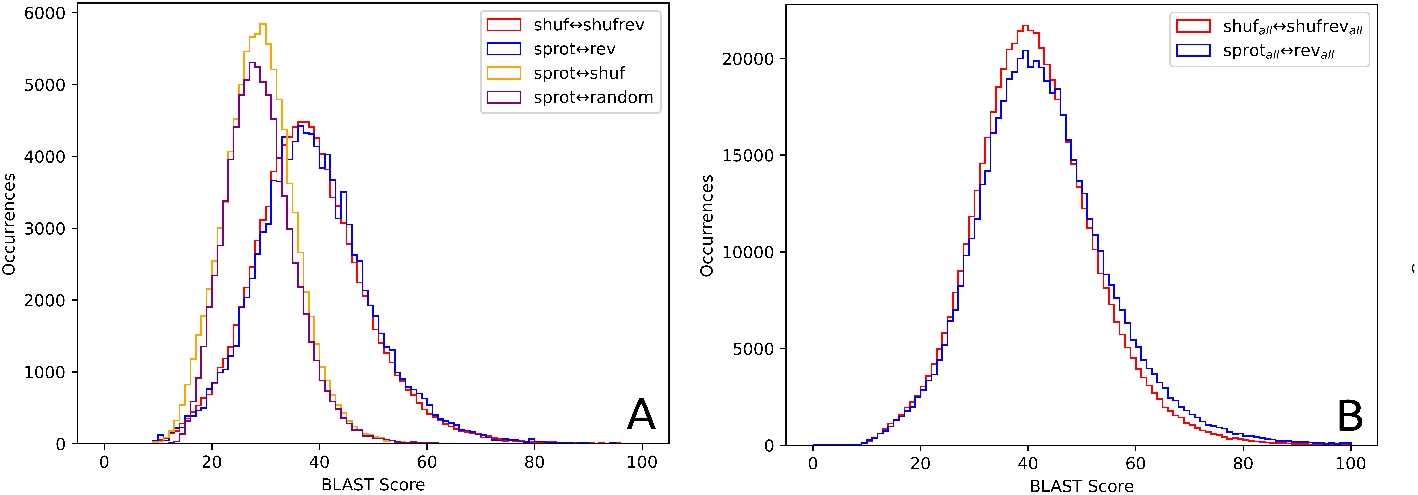
Score distributions from BLAST alignments. (A) All four pairings **sprot**↔**random, sprot**↔**shuf, sprot**↔**rev**, and **shuf**↔**shufrev** are restricted to sequences with 95% identity to SCOP2 superfamily representatives. (B) BLAST alignment scores are presented for the unfiltered sets **sprot**_**all**_↔**rev**_**all**_ and **shuf**_**all**_↔**shufrev**_**all**_.

The next two pairings involve reversed sequences. The **sprot**↔**rev** pairing aligns each of the 106,740 true Swiss-Prot proteins to the sequence’s reversal. Figure 3A shows that the distribution of these scores is strongly right-shifted relative to alignments involving **random** or **shuf**. According to common wisdom this is expected, as the argument for using reversed sequences as decoys is that real biological sequences contain patterns of repetition and regions of low complexity, and these patterns are responsible for a higher observed rate of matches against reversed decoys. This result is contrasted with the pairing **shuf**↔**shufrev**. In this pairing, each query sequence is a shuffled sequence from Swiss-Prot, and the target for each shuffled query is its reversal. The shuffled sequences retain only random levels of repeats and local low complexity, so by conventional wisdom are expected to score similarly to **sprot**↔**shuf**. However, the score distribution of **shuf**↔**shufrev** is nearly the same as that of the biological sequence and reversal, **sprot**↔**rev**. This indicates that the observed right-shift of **sprot rev** scores is primarily due to the reversal itself, and not the biological or repetitive nature of the sequences. Figure 3B presents score distributions for the reversal alignments of the unfiltered set of proteins (**sprot**_**all**_), and shows a very small right shift for **sprot**_**all**_↔**rev**_**all**_ relative to **shuf**↔**shufrev**. This shift represents the impact of repetition and low complexity, and is much smaller than the shift between **shuf**↔**shufrev** and **sprot**↔**shuf**.

Figure 4 presents scores for these various pairings as a function of length. Specifically, every pair was binned according to sequence length, then the mean *blastp* score was computed for each bin. Sequences aligned to other **sprot** sequences (**sprot**↔**random**) may differ in length, so were binned by the geometric mean length of the query and target sequences; they show a score distribution that is essentially identical to that of sequences aligned to their shuffled match (**sprot**↔**shuf**). When **sprot** sequences are aligned to their reversals (**sprot**↔**rev**), they produce consistently higher scores than they do when aligned to their shuffled partners (**sprot**↔**shuf**). The majority of the difference is explained by the simple fact of reversal, as evidenced by the gap between **sprot**↔**shuf** and **shuf**↔**shufrev** and the overlap between **sprot**↔**rev** and **shuf**↔**shufrev**.

**Fig. 4.**
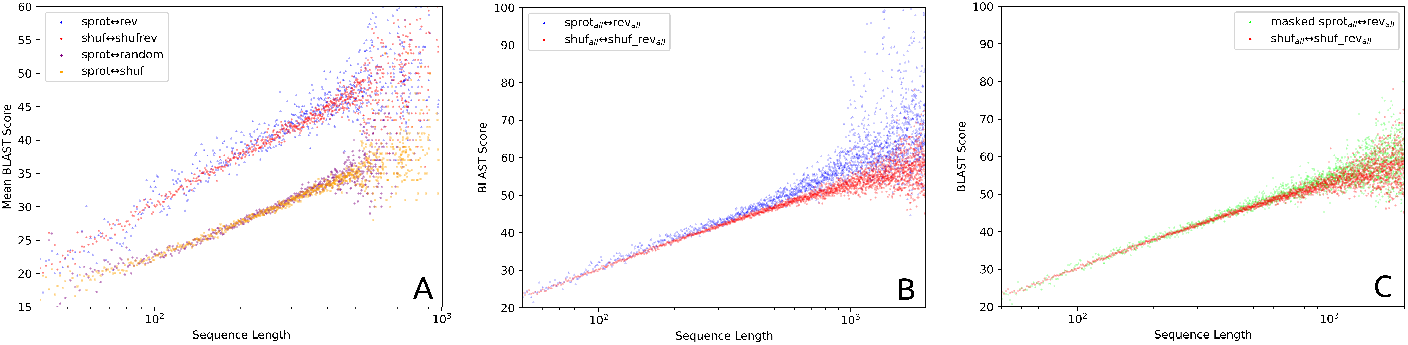
BLAST alignment scores as a function of sequence length, for the set of 106,740 sequences with full-length matches to SCOP sequences. Sequences were binned according to sequence length, and mean blastp score was computed for each bin. (A) The set **sprot**↔**random** includes pairs that may not have equal length, so scores are plotted by the geometric mean length of query and target sequence. For all remaining sets both sequences in each pair share the same length: **sprot**↔**shuf, sprot**↔**rev**, and **shuf**↔**shufrev** are plotted. (B) This plot shows scores as a function of sequence length for all 565,854 Swiss-Prot sequences with length under 2000: **sprot**_**all**_ ↔**rev**_**all**_, and **shuf**_**all**_ ↔**shufrev**_**all**_. (C) The values from B are reproduced, and supplemented with the results of masking with tantan prior to search (**masked sprot**_**all**_ ↔**rev**_**all**_); the resulting distribtuon shows that most of the modest difference between **sprot**_**all**_ ↔**rev**_**all**_ and **shuf**_**all**_ ↔**shufrev**_**all**_ is resolved by application of tandem repeat masking.

### Role of Tandem Repeats in BLAST Score Gap

In Figure 4, Swiss-Prot sequences aligned to their reversals show a modest increase in BLAST score relative to shuffled sequences to their reversals. To explore the role of tandem repeats in this shift, we created the dataset **masked sprot**_**all**_ by hard-masking repetitive regions using *tantan* (11) (see Methods). Figure 4B presents these analyses, focused on **shuf**_**all**_↔**shufrev**_**all**_ and **sprot**_**all**_↔**rev**_**all**_ distributions, while Figure 4C replaces **sprot**_**all**_↔**rev**_**all**_ with another distribution showing the result of first masking with *tantan* (11) (default settings), then searching against its reversal (**masked sprot**_**all**_↔**rev**_**all**_). These results demonstrate that the small gap between **shuf**_**all**_↔**shufrev**_**all**_ and **sprot**_**all**_↔**rev**_**all**_ is almost entirely due to repetitive sequence that can be effectively masked. This phenomenon, and its response to masking, reoccurs in the analyses replicated on DNA sequences in the corresponding section below.

### The Role of Mutations (Approximate Reversals)

The above methods have only examined alignments between a sequence and its exact reversal. In practice, a database of reversed se-quences may contain the reversal of a homolog to the query sequence, rather than the query itself. It seems reasonable to expect the effect of reversal on alignment score to be diminished, but perhaps not entirely removed. To simulate this situation, we produced a collection of reversal variants by sampling amino acid substitution mutations from conditional distributions with self-emission probabilities ranging from 46% to 90% (see Methods). These transformations are of **shufrev**_**all**_, named **shufrev**_**all**_**-46pct, shufrev**_**all**_**-50pct, shufrev**_**all**_**-60pct, shufrev**_**all**_**-70pct, shufrev**_**all**_**-80pct**, and **shufrev**_**all**_**-90pct**. Pairwise alignments are performed for each transform to **shuf**_**all**_ using the Smith-Waterman (SW) algorithm implemented in *BioAlignments*.*jl* (30). SW was used instead of blastp in order to maximize the number of alignment score datapoints captured, because blast often produces no alignment (or score) for a comparison between a query and decoy.

The SW alignments between **shuf**_**all**_ and **shufrev**_**all**_**-Xpct** for X ∈{46, 50, 60, 70, 80, 90} were computed using the BLOSUM62 scoring matrix and gap penalties set to −11 for opening and −1 for extension, to mimic the default configuration of *blastp*. Figure 5 displays the spread of these scores, as well as those of **sprot**_**all**_↔**shuf**_**all**_ and **shuf**_**all**_↔**shufrev**_**all**_. The self-matching effect of reversal is lost completely in **shufrev**_**all**_**-46pct** and almost undetectable in **shufrev**_**all**_**-50pct**, which present score distributions similar to **sprot**_**all**_↔**shuf**_**all**_. But when simulated homologs are created with higher average percent identity between query and partner, and the partner is then reversed (yielding **shufrev**_**all**_**-Xpct** for X ∈{60,70,80,90}), the associated pairs exhibit higher-than-random affinity between sequences and their (approximate) reversal. Thus, there is a risk of overstating false match rates even when the reversed decoy set contains homologs sharing 60% or greater similarity to query sequences.

**Fig. 5.**
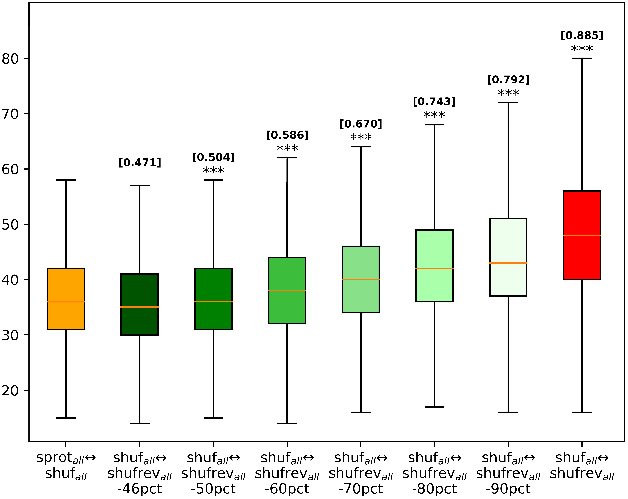
Box-and-whisker plot of SW alignment score distributions for alignment pairs **sprot**_**all**_ ↔**shuf**_**all**_, **shuf**_**all**_ ↔**shufrev**_**all**_, and **shufrev**_**all**_**-Xpct** for X ∈ {46, 50, 60, 70, 80, 90}. Above each box plot, the symbol *** indicates distributions that are significantly higher than **sprot**_**all**_ ↔**shuf**_**all**_ based on the Wilcoxon rank test (P = 0, due to computational underflow). The values [Y] indicates the fraction of shuffled instances for which the associated box plot produces a higher score than **sprot**_**all**_ ↔**shuf**_**all**_.

### Sequences match their own reversals much more than other reversals

The previous results show that alignment of a shuffled sequence to its reversal produces a score that is, on average, roughly 50% higher than alignment of that shuffled sequence to the original protein. In the context of high scoring random matches, this mean score is less important than the frequency of very high scoring outliers. To explore the right tail of this score distribution, we captured a slice of all sequences in **sprot** with length between 280 and 320, and computed the frequency with which scores are observed in (i) alignments of each sequence in this set with a shuffled copy of the sequence, (ii) alignments of shuffled copies of each sequence in this set with reversed copies of those shuffled copies, and (iii) alignments of shuffled copies of each sequence in this set with reversed copies that we mutated according to the 80pct substitution matrix from the previous section.

Figure 6 shows that alignment to reversals produces a highly right-skewed distribution, and that this right skew is strongly evident even when reversals have been subjected to mutation to 80% identity. All three score distributions appear to be well modeled by a Gumbel distribution, so we fit Gumbel parameters using the maximum likelihood estimation fitting function from Distributions.jl (32). As a concrete example, a score of 165 is expected to occur by chance with a probability of ∼3e-12 in masked sequence-to-shuffled analysis. Meanwhile the same score is expected to occur with frequency ∼4.5e-7 for shuffled sequence aligned to *reversed* masked sequences.

**Fig. 6.**
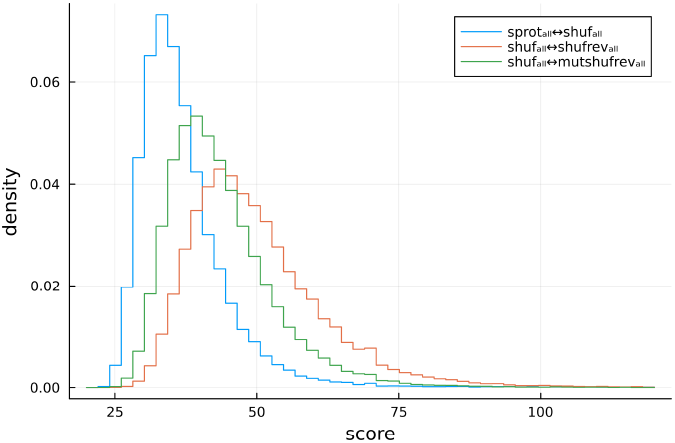
Observed score frequencies for three types of aligned pairs, using only sequences of length 280-320. The three pairings are: (i) sequences aligned to shuffled copies (blue), (ii) those shuffled copies aligned to their reversals (orange), and (iii) the shuffled copies aligned to their reversals that have been subjected to mutation resulting in 20% amino acid change (green).

To further explore the overabundance of high scores in self-reversal alignments, we aligned a set of 10,000 sequences against reversals of that set, and found that sequences were much more likely to find a high-scoring match to their own reversals than to the reversals of any other sequences. Specifically, we selected 10,000 random sequences from UniProtKB/TrEMBL (33) and masked tandemly repetitive regions in each sequence using tantan (11) with default settings, resulting in a set that we call. Each sequence in this set was reversed, resulting a set that we call. Blast was used to search against the database, so that each query sequence in was aligned to the 10,000 reversed sequences in. Under conventional reasoning, the reversed sequences in are unrelated to the original sequences in, so each query should have a similar chance of matching any of the sequences in ; thus, a query sequence should produce a high-scoring alignment to the reversal of some *other* sequence roughly 10,000x as often as it will produce a high-scoring alignment to its own reversal. In conflict with this expectation, among all blastp matches with E-value *<* 0.001, there were 12 matches between any query and its self reversal, and only 3 matches between a query and any of the 9,999 other reversals.

We highlight that these self-reversal matches are not due to repetitive sequence, since repeat sequences were masked before alignment. Because low complexity (biased use of the available protein alphabet) could cause elevated rate of self-reversal matches, we assessed sequence complexity; since blastp does not report a measure of sequence complexity, we also performed alignment using phmmer (25), and found that matches reported negligible composition bias.

### Exact palindromes in Swiss-Prot

To gain insight into the high frequency of high-scoring alignments between sequences and their reversals, we computed all maximal-length palindromes in each of the 565,855 sequences in **sprot**_**all**_ as well as **shuf**_**all**_ for a control, using our implementation of Manacher’s algorithm (31). An additional control was observed by computing the longest common substring (LCS) between sequence pairs in **sprot**_**all**_ and **shuf**_**all**_.

The palindromic substrings in **sprot**_**all**_, **shuf**_**all**_, and the LCS between them, were binned by the length of the sequence containing the substring; averages computed per each length are presented in Figure 7. The length distribution of maximal palindromes is closely related to that of longest common substrings; a result of Duchon et al. (20) implies that the expected length of (ungapped) maximal palindromes is asymptotically equivalent to the expected length of longest common substrings as derived by Arratia et al. (19). In order to contextualize our analyses within this theory, we plotted the expected lengths of (i) LCS between two random sequences 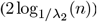, and (ii) palindrome within a sequence 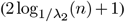.

**Fig. 7.**
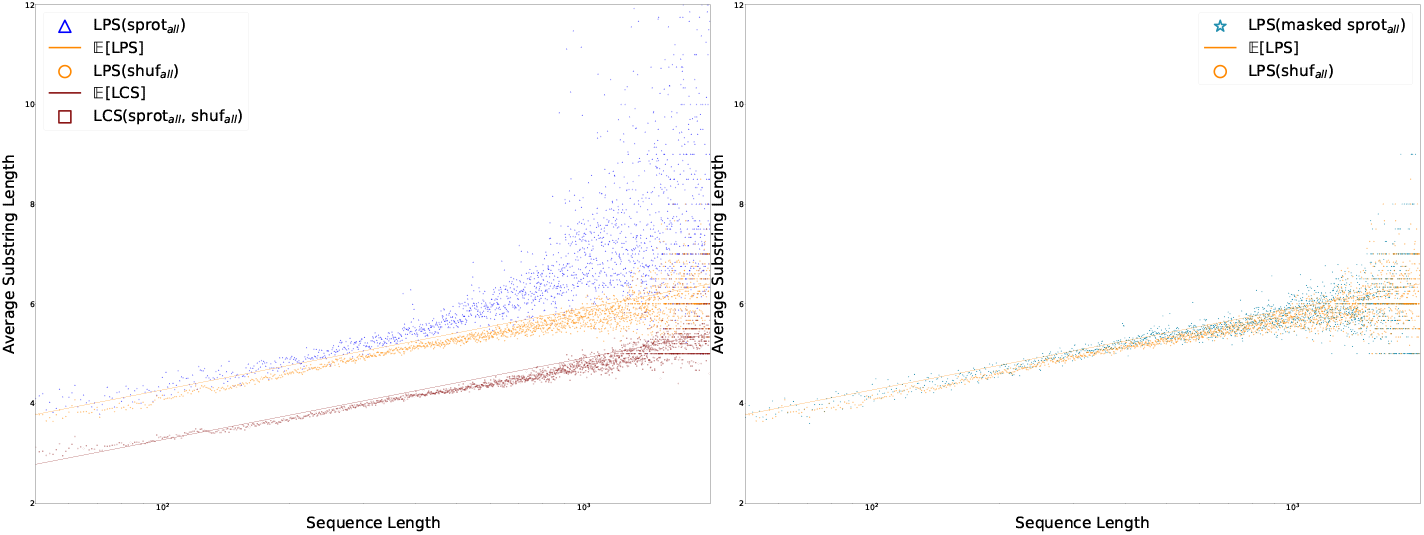
Lengths of maximal palindromes occurring in biological sequences **sprot** and random sequences **shuf** with the expected lengths derived from (20). Biological palindromes appear to be asymptotically longer than random palindromes. On the left, the lengths of longest common substrings (LCS) of pairs between **sprot** and **shuf** are plotted with the expected length from (19). On the right, maximal palindromes remaining after masking **sprot** with *tantan* are plotted; with repetitive regions removed, the lengths of biological palindromes become closely clustered around the random expectation.

Figure 7 shows that **sprot**_**all**_ contains, on average, longer palindromic substrings than **shuf**_**all**_. While palindromes found in the shuffled dataset are densely concentrated around their expected value, the average palindrome length in **sprot**_**all**_ diverges from the expectation, particularly as sequence length increases. This asymptotic behavior only yields a large divergence from expected score in the small fraction of sequences in **sprot**_**all**_ that are longer than ∼500 amino acids.

To explore the role of repetitive sequence in this shift, we computed maximal palindromes for each *masked* sequence. The resulting distribution of palindrome lengths (Figure 7B) mirrors that of the shuffled sequences in Figure 4C, clustering densely around the expected value. Based on manual review, the divergence of biological palindromes from their stochastic counterparts appears to be almost entirely explained entirely by long tandem repeats. In particular, these palindromes are composed of smaller, palindrome-supporting sub-units like ‘PNAN’ or ‘SD’, leading respectively to palindromes like PNANPNANPNANP and SDSDSDSDS.

### Palindromes in the context of mass spectrometry

When using mass spectrometry to identify proteins in a sample, a library of reversed sequences is usually used to set thresholds based on risk of false labeling. (34) Here, we consider the potential that including reversals in mass spectrometry benchmarks may lead to overestimated false match rates and a resulting decrease in search sensitivity.

With all sequences in **sprot**_**all**_, we produced protein fragments similar to those used in mass spectrometry, splitting each sequence at each instance of a lysine or arginine, except where followed by a proline (see Methods). After splitting, substrings with length between 5 and 100 were accumulated into sets named **sprot**_**tr**_. Fragments in **sprot**_**tr**_ were shuffled to produce a related set **shuf**_**tr**_.

Figure 8 displays the average lengths of longest palindromic subsequence (LPS) in each set of strings, binned by the length of the string. To place these in context, the longest common substring (LCS) was computed between paired sequences in **sprot**_**tr**_ and **shuf**_**tr**_.

**Fig. 8.**
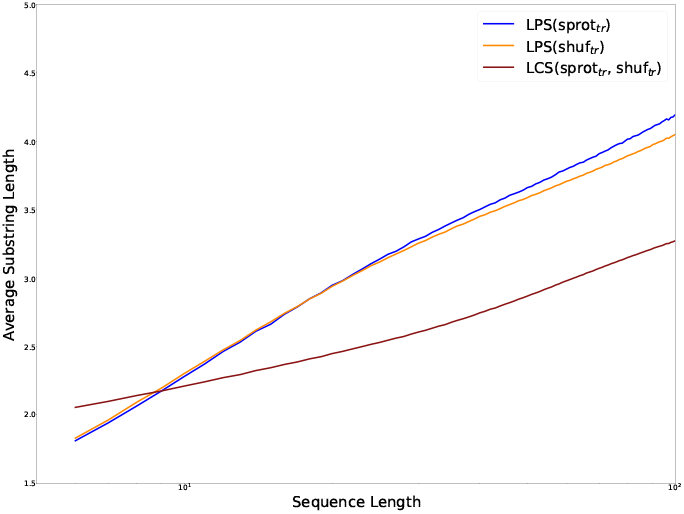
The average length of LPS in tryptic peptides between lengths 5 and 100, generated from an in-silico trypsin digest of **sprot**_**all**_, are plotted against those of tryptic substrings occurring **shuf**_**all**_. When tantan-masked regions are removed as in **masked sprot**_**all**_, the difference in distributions of LPS length collapses.

Trends established by the preceding results are still apparent in Figure 8. LPS are longer than LCS, and because tryptic peptides are so short, the constant component separating LPS and LCS lengths is large relative to their shared logarithmic component. However, the relationship between LPS and LCS appears to invert for very short sequences. The data indicate that *LCS*(**sprot**_**tr**_, **shuf**_**tr**_) *> LP S*(**sprot**_**tr**_) for se-quences shorter than 10 amino acids, and *LP S*(**sprot**_**tr**_) ∼ *LP S*(**shuf**_**tr**_) for sequences shorter than approximately 20 peptides. Composition bias in short tryptic peptides may contribute to quasi-invariance under shuffling.

### Palindromes in DNA

The analyses presented above have been focused on protein sequences, but the surprising prevalence of palindromes is a general feature of strings, regardless of alphabet. As with the protein results shown in this manuscript, a random DNA sequence (i.e. one with no repetitive or low-complexity regions) will on average have a longer exact match to its reversal than it will to a shuffled version of itself; it will also typically produce a higher score when aligned (under a reasonable scoring scheme) to its reversal than when aligned to a shuffled version of itself.

To explore this, we analyzed DNA occurring in human chromosome 22. Chromosome 22 was downloaded and annotated using the UCSC Genome Browser. Gene sequences were extracted from the chromosome using GffRead; introns and intergenic regions were also sampled from chromosome 22 to match the length distribution of genes. These datasets are respectively denoted **mRNA, intron**, and **intergenic**. Each set was filtered to contain sequences less than 50000 bp. As above, variant sets were created using shuffling and masking, respectively denoted **shuf**_**mRNA**_, **shuf**_**intron**_, **shuf**_**intergenic**_, and **masked mRNA, masked intron**, and **masked intergenic**. For more detail, see the preceding section in Methods.

The LPS of each DNA sequence was computed using Manacher’s algorithm. LCS were constructed between each se-quence and a shuffled counterpart using a substring automata. Fig 9 plots the lengths of these common substrings, binned by sequence length.

**Fig. 9.**
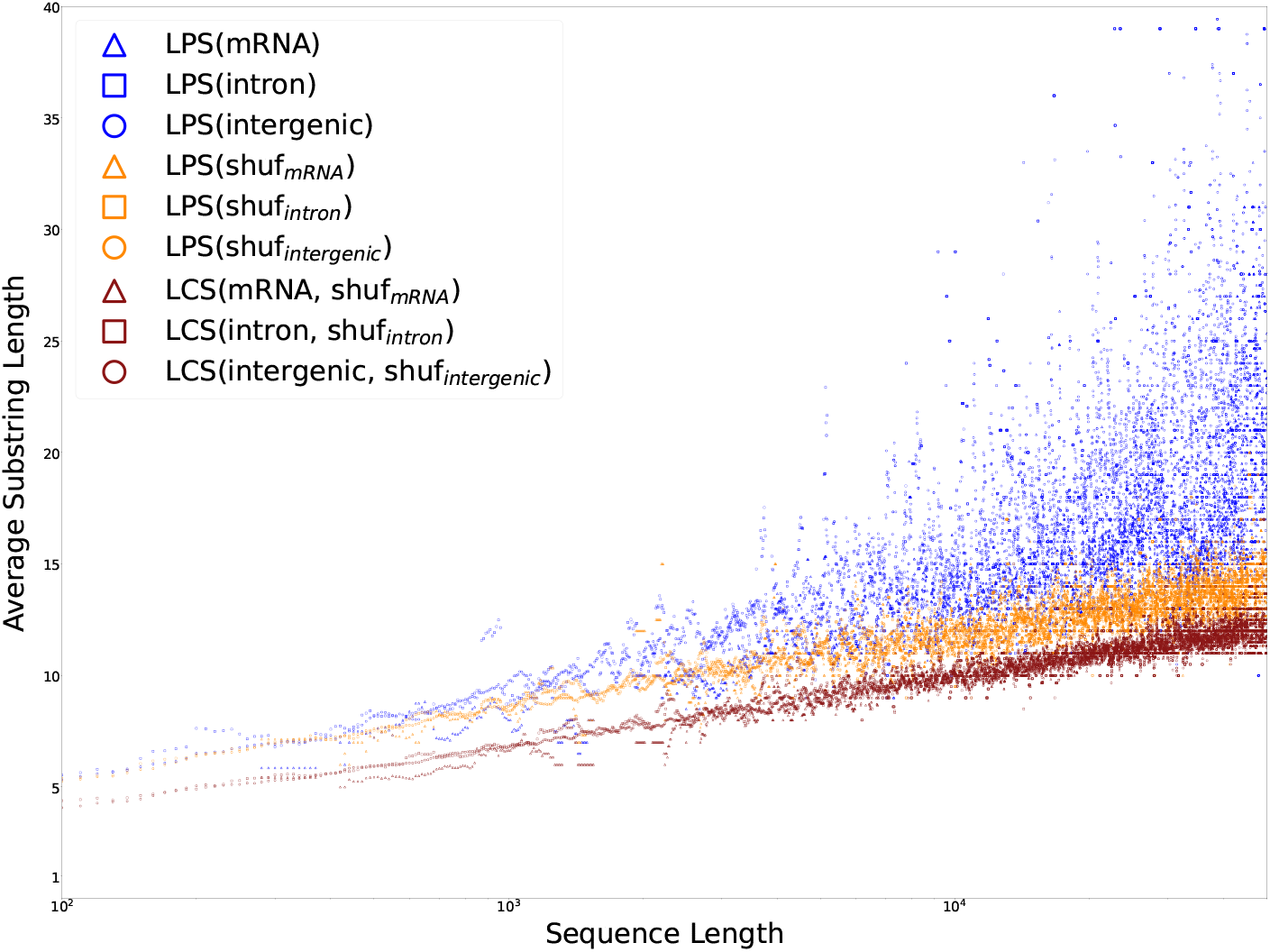
The average lengths of palindromes occurring in human chromosome 22 are plotted by sequence length against shuffled controls.

**Fig. 10.**
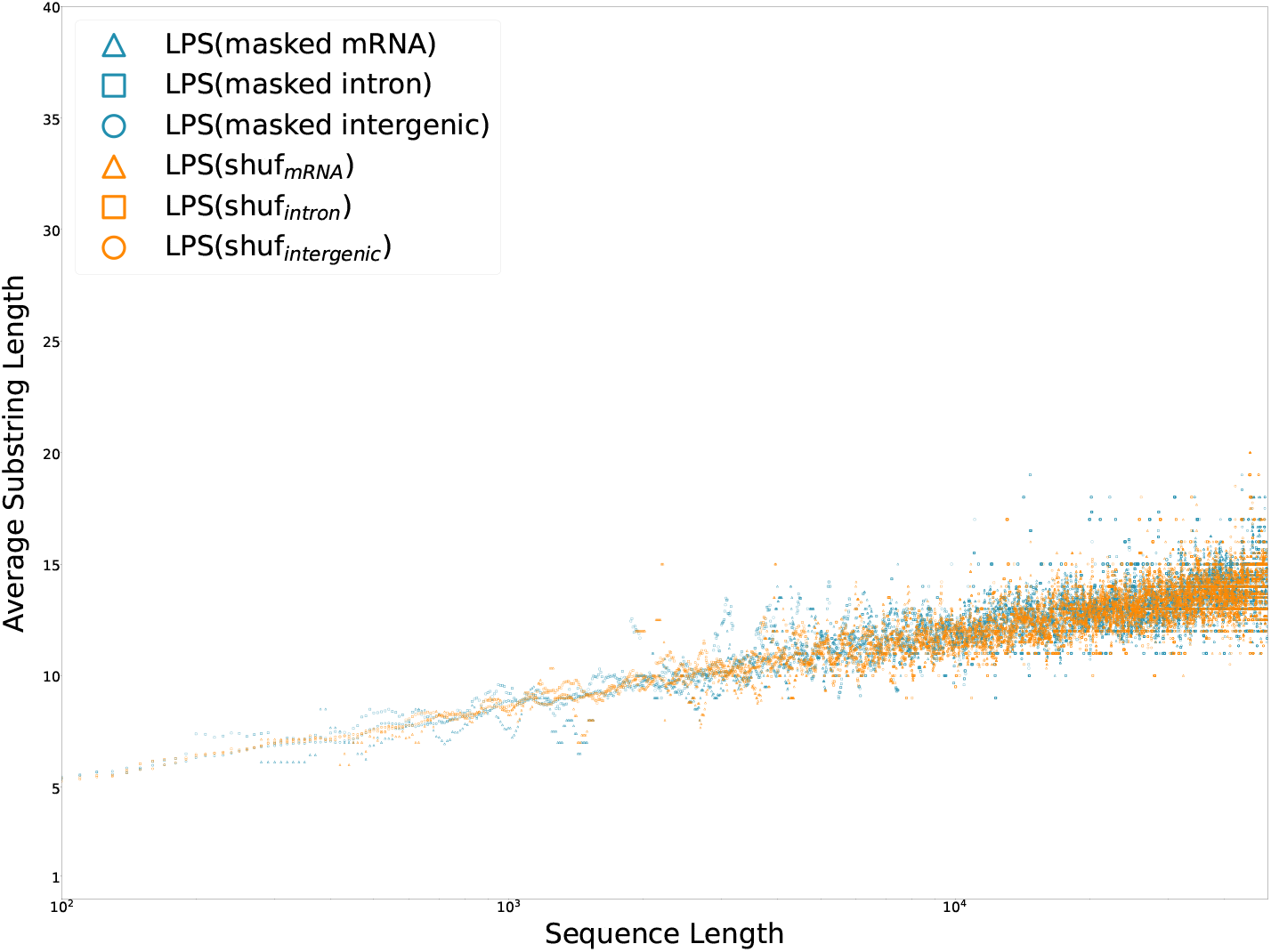
Repetitive regions in human chromosome 22 are masked via tantan; the remaining palindromes in non-repetitive DNA, as well as those in corresponding shuffled sequences, are plotted. Both data are distributed closely around their expectation. As above, when repetitive subsequences are suppressed, the biological contribution to reverse affinity appears negligible.

The analysis of DNA reproduces the results observed in peptides and proteins. On average, the longest palindromic sub-string of a sequence is longer than the longest common sub-string between that sequence and a random other. When biologially-induced repetitions are present, the difference between LPS and LCS grows superlogarithmically. In the absence of repetition (either by shuffling or tantan masks), this difference appears constant.

### Summary of Results

In summary, alignment to reversal produces a substantial increase in score compared to alignment to unrelated sequences, even when there is no inherent repetitiveness or low complexity in the initial sequence. This effect is diminished when the reversed sequence contains mutations relative to the query, but remains significant for related sequences exceeding 60% identity to the non-reversed sequence. As a result, when reversed biological sequences are used as benchmarks they may overstate the risk of false matches.

## Discussion

Up to this point, we have discussed the prevalence of palindromes in terms of their impact on bioinformatic analysis (specifically, overstatement of false positive rates in sequence search). It is also interesting to consider how this existence of palindromes in sequences may play a role in biological processes. The role of palindromes in proteins is currently unknown and research on their function is ongoing (35–38). Sheari et al. (36) argue that the prevalence of palindromic regions in proteins is a side-effect of repetitive and low-complexity regions. Sridhar et al. (37) found that 60% of palindromes in proteins are “not associated with any regular secondary structure”, consistent with the notion that palindromes in proteins are not dependent on repetitiveness such as that observed in alpha helical structures. Furthermore, 80% of palindromes in proteins do not interact with active sites, catalytic residues, ligands, or metal ions. In a more direct study, Aikaterini et al. (38) synthesized a 4-alpha-helical bundle and its reversal, and demonstrated that while the reversed sequence does fold into a helix – likely due to the aforementioned effects of repetition in helical sequences – the resulting structure was not a true alpha helix. In general, these results are consistent with our results above, in that the frequency of palindromes in true proteins appears to be primarily a function of the frequency of palindromes in random sequences, and repetitive aspects of some sequences may lead to a moderate increase in the observed rate of palindromes.

### A better benchmark?

Can we compensate for the over-abundance of palindromic regions in biosequence benchmarks? The supplemental materials include a script for generating improved reversed decoy benchmarks, given a query set *Q* and a target set *T*. The target set *T* is restricted to *T* ^′^, containing sequences with less than 50% percent identity to any sequence in *Q*. The reversal of *T* ^′^, denoted *R*, retains the original biological signal present in *T* and omits reversal-induced false matches. As above, it may be necessary to mask repetitive regions before performing the initial alignment.

## Author contributions statement

TJW conceived the experiment(s). GGH wrote all software, conducted experiments and and developed analysis of expected palindrome lengths. GGH and TJW analysed the results and wrote and reviewed the manuscript.

## ACKNOWLEDGEMENTS

We wish to thank Dave Rich for careful consideration of benchmarking issues during his thesis work, and for discussions that motivated early work on this analysis. We also thank Robert Hubley and Daniel Olson for illuminating conversations regarding both palindrome theory and application to annotation benchmarks. We also thank Martin Frith for providing valuable feedback and insights in response to our first preprint draft. Analyses were made possible thanks to high performance computing (HPC) resources supported by the University of Arizona TRIF, UITS, and Research, Innovation, and Impact (RII) and maintained by the UArizona Research Technologies department. GGH and TJW were supported by NSF DBI 1933305.

